# Enhancing senior high school student engagement and academic performance using an inclusive and scalable inquiry-based program

**DOI:** 10.1101/822783

**Authors:** Locke Davenport Huyer, Neal I. Callaghan, Sara Dicks, Edward Scherer, Andrey I. Shukalyuk, Margaret Jou, Dawn M. Kilkenny

**Affiliations:** Institute of Biomaterials and Biomedical Engineering, University of Toronto, Toronto, Ontario, Canada; Chemical Engineering and Applied Chemistry, University of Toronto, Toronto, Ontario, Canada; Translational Biology and Engineering Program, Ted Rogers Centre for Heart Research, University of Toronto, Toronto, Ontario, Canada; George Harvey Collegiate Institute, Toronto District School Board, Toronto, Ontario, Canada

## Abstract

The multi-disciplinary nature of science, technology, engineering and math (STEM) careers often renders difficulty for high school students navigating from classroom knowledge to post-secondary pursuits. Discrepancies between the knowledge-based high school learning approach and the experiential approach of undergraduate studies leaves some students disillusioned by STEM. We present *Discovery*, a semester-long inquiry-focused learning model delivered by STEM graduate students in collaboration with high school educators, in the context of biomedical engineering. Entire classes of high school STEM students representing diverse cultural and socioeconomic backgrounds engaged in iterative, problem-based learning designed to emphasize critical thinking concomitantly within the secondary school and university environments. Assessment of grades and survey data suggested positive impact of this learning model on students’ STEM pursuits, notably in under-performing cohorts, as well as repeating cohorts that engage in the program on more than one occasion. *Discovery* presents a scalable platform blurring the divide between secondary and post-secondary learning, providing valuable learning opportunities and capturing cohorts of students that might otherwise be under-engaged in STEM.

## 1 Introduction

High school students with diverse STEM interests often struggle to understand the STEM experience outside the classroom^1^. The multi-disciplinary nature of many careers fosters a challenge for many students when considering the transition between high school study and future academic pursuits. Furthermore, this challenge is amplified by the known discrepancy between the knowledge-based learning approach common in high schools and the experiential, mastery-based approaches afforded by the undergraduate model^2^. In the latter, focused classes, interdisciplinary concepts, and laboratory experiences allow for the application of accumulated knowledge, practice in problem solving, and development of both general and technical skills^3^. Such immersive cooperative learning environments are difficult to establish in the secondary school setting and many high school educators struggle to implement within their classroom^4^. As such, high school students may become disillusioned before graduation and never experience an enriched learning environment, despite their inherent interests in STEM^5^.

Early introduction to varied math and science disciplines throughout high school is vital if students are to pursue STEM fields, especially within engineering^6^. In the context of STEM education and career choices, student self-efficacy regarding research skills has been shown to predict undergraduate student aspirations for research careers^7^. Self-efficacy has also been identified to influence ‘motivation, persistence, and determination’ in overcoming challenges in a career pathway^8^. It is suggested that high school students, when given opportunity and support, are capable of successfully completing rigorous programs at STEM focused schools^9^. However, alternate studies have shown no significant differences in participation rates in advanced sciences and mathematics for these students compared to their peers at non-STEM focused schools^10^. In fact, Brown *et al* studied the relationships between STEM curriculum and student attitudes, and found the latter played a more important role in intention to persist in STEM when compared to self-efficacy^11^. Therefore, creation and delivery of modern and exciting curriculum is fundamental to engage and retain students.

Many public institutions support the idea that post-secondary led engineering education programs are effective ways to expose high school students to engineering education and relevant career options, and also increase engineering awareness^12^. Although singular class field trips are used extensively to accomplish such programs, these may not allow immersive experiences for application of knowledge and practice of skills that are proven to impact long-term learning and influence career choices^13,14^. Longer-term immersive research experiences, such as after-school programs or summer camps, have shown successful at recruiting students into STEM degree programs and careers, where longevity of experience helps foster self-determination and interest-led, inquiry-based projects^3,7,15-17^. Such activities convey the elements that are suggested to make a post-secondary led high school education program successful: hands-on experience, self-motivated learning, real-life application, immediate feedback, and problem-based projects^18,19^. In combination with immersion in university teaching facilities, learning is authentic and relevant, and consequently representative of an experience found in actual STEM practice^20^.

Supported by the outcomes of previously identified effective program strategies, University of Toronto (U of T) graduate trainees created *Discovery*, a novel high school education program, to develop a comfortable yet stimulating environment of inquiry-focused iterative learning for senior high school students. Built in strong collaboration with science educators from George Harvey Collegiate Institute (Toronto District School Board), *Discovery* stimulates application of STEM concepts within a unique semester-long applied curriculum delivered iteratively within both U of T undergraduate teaching facilities and collaborating high school classrooms^21^. Based on the volume of medically-themed news and entertainment that is communicated to the population at large, the rapidly-growing and diverse field of biomedical engineering (BME) was considered an ideal program context^22^. In its definition, BME necessitates cross-disciplinary STEM knowledge focused on the betterment of human health, wherein *Discovery* facilitates broadening student perspective through engaging inquiry-based projects. Importantly, *Discovery* allows all students within a class cohort to work together with their classroom educator, stimulating continued development of a relevant learning community that is deemed essential for meaningful context and important for transforming student perspectives and understandings^23,24^. Multiple studies support the concept that relevant learning communities improve student attitudes towards learning, significantly increasing student motivation in STEM courses, and consequently improving the overall learning experience^25^. Learning communities, such as that provided by *Discovery*, also promote the formation of self-supporting groups, greater active involvement in class, and higher persistence rates for participating students^26^.

The objective of *Discovery*, through structure and dissemination, is to engage senior high school science students in challenging, inquiry-based practical BME activities as a mechanism to stimulate comprehension of STEM curriculum application to real world concepts. Consequent focus is placed on critical thinking skill development through an atmosphere of perseverance in ambiguity, something not common in a secondary school knowledge focused delivery but highly relevant in post-secondary STEM education strategies. Herein, we describe the observed impact of the differential project-based learning environment of *Discovery* on student performance and engagement. We specifically hypothesize that value of an inquiry-focused model is tangible for students that struggle in a knowledge focused delivery structure, where engagement in conceptual critical thinking in the relevant subject area stimulates student interest and resulting academic performance. Assessment of these outcomes suggests that when provided with a differential learning opportunity, the performance of these students increased as they engaged more thoroughly in STEM subject matter. Consequently, *Discovery* provides a framework to the potential efficacy of the model for scalable application in bridging the gap in critical thinking and problem solving between secondary and post-secondary education.

## 2 Results

### 2.1 Program Delivery

The outcomes of the current study result from execution of *Discovery* over five independent academic terms as a collaboration between IBBME (graduate students, faculty, and support staff) and George Harvey Collegiate Institute (science educators and administration) stakeholders. Each term, the program allowed senior secondary STEM students (Grades 11 and 12) opportunity to engage in a novel project-based learning environment. The program structure uses the engineering capstone framework as a tool of inquiry-focused learning objectives, motivated by a central BME global research topic, with research questions that are inter-related but specific to the curriculum of each STEM course subject (**Fig 1**). Over each 12-week term, students worked in teams (3-4 students) within their class cohorts to execute projects with the guidance of U of T trainees (*Discovery* instructors) and their own high school educator(s). Student experimental work was conducted in U of T teaching facilities relevant to the research study of interest (i.e., Biology and Chemistry-based projects executed within Undergraduate Teaching Laboratories; Physics projects executed within Undergraduate Design Studios). Students were introduced to relevant techniques and safety procedures in advance of iterative experimentation. Importantly, this experience served as a course term project for students, who were assessed at several points throughout the program for performance in an inquiry-focused environment as well as within the regular classroom (**Fig 1; S3 Appendix III**). To instill the atmosphere of STEM, student teams delivered their outcomes in research poster format at a final symposium, sharing their results and recommendations with other post-secondary students, faculty, and community in an open environment. An exemplary term of student programming can be found in **S1 Appendix I**.

**Fig 1.**
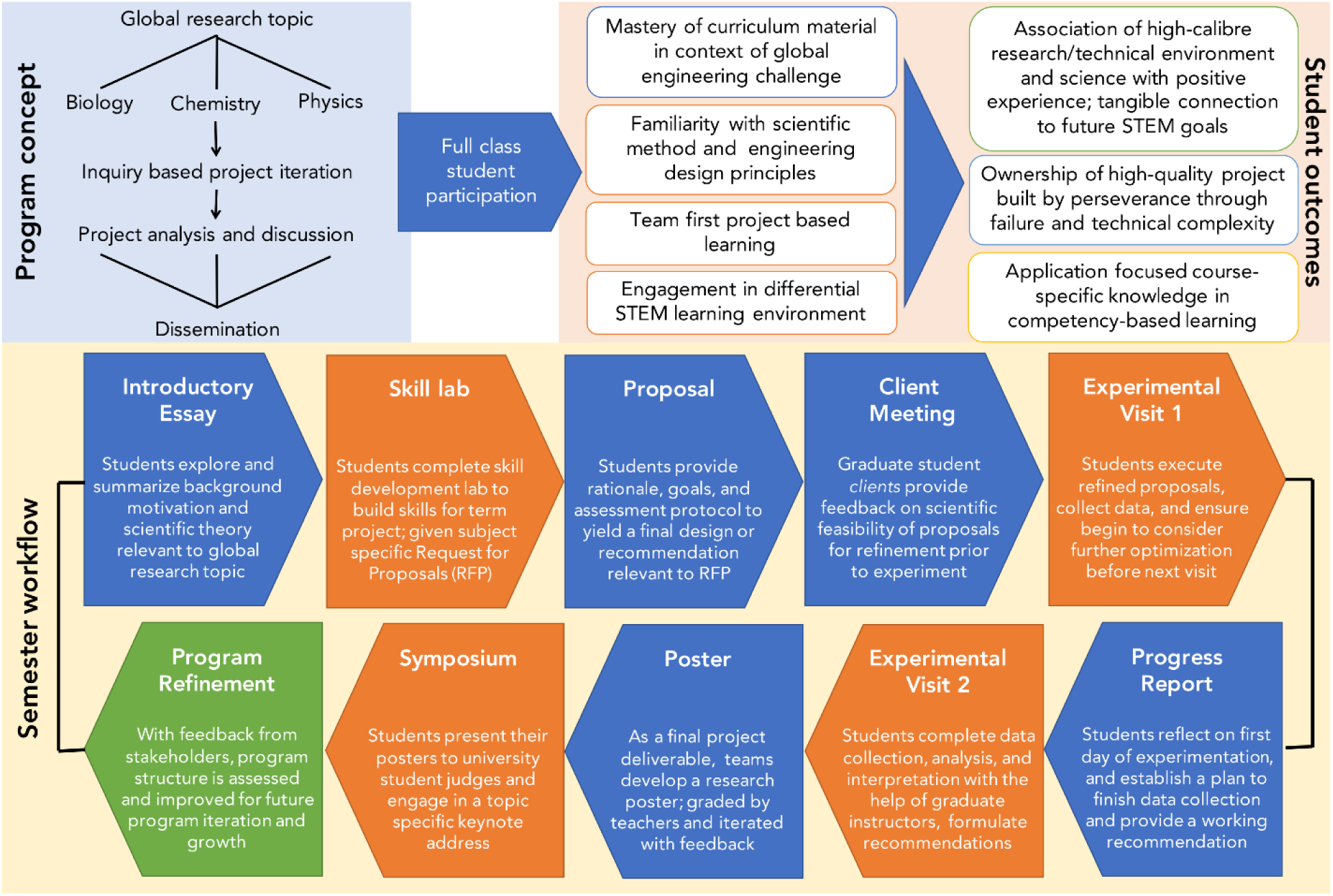
Structure and rationale underlying the *Discovery* framework. The general program concept (blue background; *top left*) highlights a global research topic examined through student dissemination of subject specific research questions, yielding multifaceted student outcomes (orange background; *top right*). Each program term (semester workflow, yellow background; *bottom panel*), students work on program deliverables in class (blue), iterate experimental outcomes within university facilities (orange), and are assessed accordingly at numerous deliverables in an inquiry-focused learning model (**S3 Appendix III**).

Over the course of five semesters there were 268 instances of tracked student participation, representing 170 individual students. Specifically, 94 students participated during only one semester of programming, 57 students participated in two semesters, 16 students participated in three semesters, and 3 students participated in four semesters. Multiple instances of participation represent students that enrol in more than one STEM class during their senior years of high school, or who participated in Grade 11 and subsequently Grade 12. All assessments were performed by high school educators for their respective STEM class cohorts using consistent grading rubrics and assignment structure (summarized in **S3 Appendix III**). Here, we discuss the outcomes of student involvement in this experiential curriculum model.

### 2.2 Student performance and engagement

Student grades were assigned, collected and anonymized by educators for each *Discovery* deliverable (background essay, client meeting, proposal, progress report, poster and final presentation). Educators anonymized collective *Discovery* grades, the component deliverable grades thereof, final course grades, attendance in class and during programming, as well as incomplete classroom assignments for comparative study purposes. Students performed significantly higher in their cumulative *Discovery* grade than in their cumulative classroom grade (final course grade less the *Discovery* contribution; p < 0.0001). Nevertheless, there was a highly significant correlation (p < 0.0001) observed between the grade representing combined *Discovery* deliverables and the final course grade (**Fig 2a**).Further examination of the full dataset revealed two student cohorts of interest: the “Exceeds Expectations” (EE) subset (defined as those students who achieved ≥ 1 SD [18.0%] grade differential in *Discovery* over their final course grade; N=99 instances); and the “Multiple Term” (MT) subset (defined as those students who participated in *Discovery* more than once; 76 individual students that collectively accounted for 174 single terms of assessment out of the 268 total student-semesters delivered) (**Fig 2b-c**). These subsets were not unrelated; 46 individual students who had multiple experiences (60.5% of total MTs) exhibited at least one occasion in achieving a ≥18.0% grade differential. MT students that participated in 3 or 4 semesters (N = 16 and 3, *respectively*) showed no significant increase by linear regression in their course grade over time (p = 0.40, **Fig 2e**), but did show a significant increase in their *Discovery* grades (p = 0.0009, **Fig 2f**).

**Fig 2.**
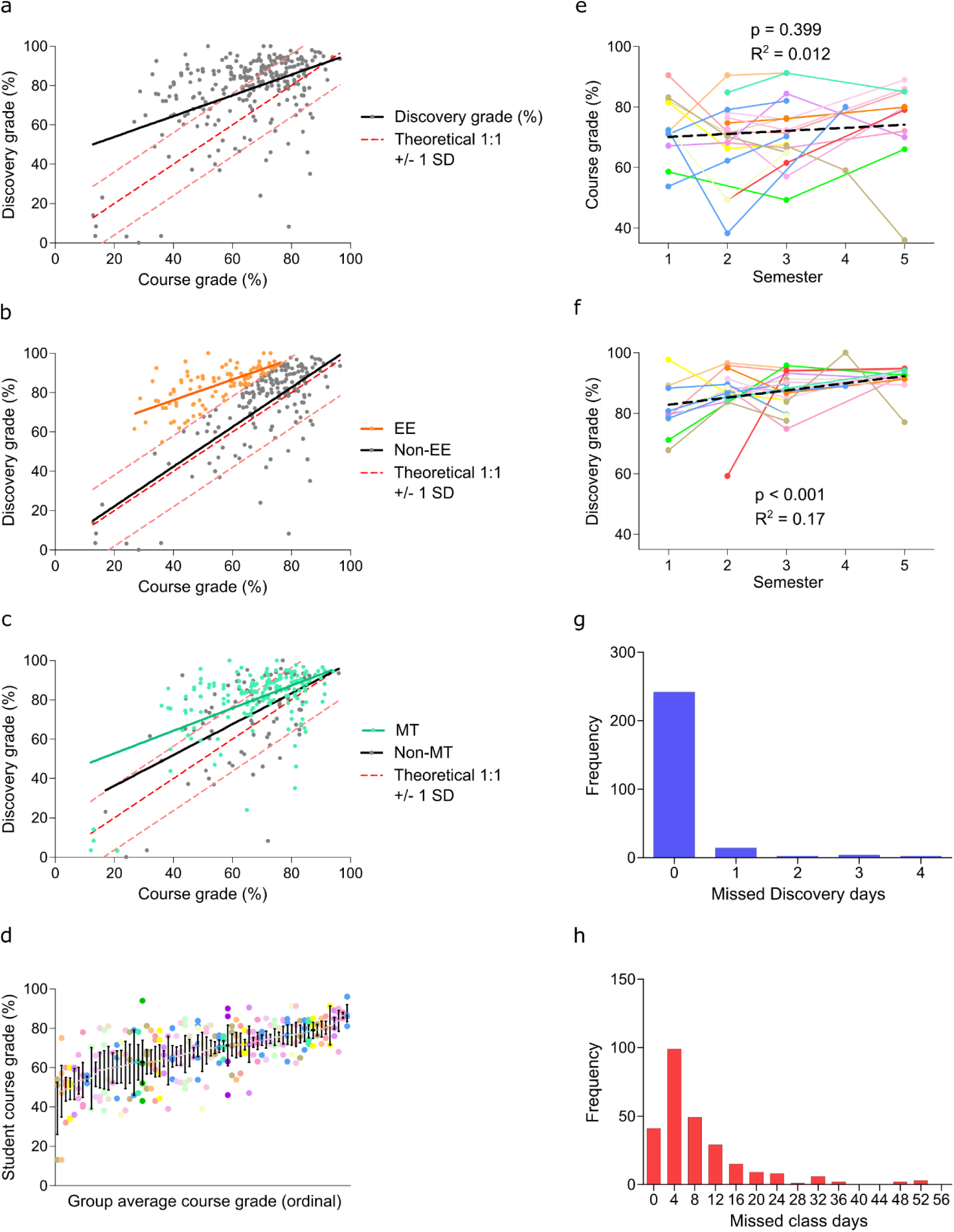
Student aggregate performance in *Discovery* and identification of subsets. (a) Linear regression of student grades reveals a significant correlation (p = 0.0009) between *Discovery* performance and final course grade less the *Discovery* contribution to grade, as assessed by educators. The dashed red line and intervals represent the theoretical 1:1 correlation between *Discovery* and course grades and standard deviation of the *Discovery*-course grade differential, respectively. (b & c) Identification of subgroups of interest, Exceeds Expectations (EE; N = 99, *orange*) who were ≥ +1 SD in *Discovery*-course grade differential and Multi-Term (MT; N = 174, *teal*), of which N = 65 students were present in both subgroups. (d) Students tended to self-assemble in working groups according to their final course performance; data presented as mean ± SEM. (e) For MT students participating at least 3 semesters in *Discovery*, there was no significant correlation between course grade and time, while (f) there was a significant correlation between *Discovery* grade and cumulative semesters in the program. (g & h) Histograms of total absences per student in (g) *Discovery* and (h) class (binned by 4 days to be equivalent in time to a single *Discovery* absence).

As students participated in group work, there was concern that lower-performing students might negatively influence the *Discovery* grade of higher-performing students (or vice versa). However, students were observed to self-organize into groups where all individuals received similar final overall course grades (**Fig 2d**), thereby alleviating these concerns. In addition, students demonstrated excellent *Discovery* attendance; at least 91% of participants attended all *Discovery* sessions in a given semester (**Fig 2g**). In contrast, class attendance rates reveal a much wider distribution where 60.8% (163 out of 268 students) missed more then 4 classes (equivalent in learning time to one *Discovery* session) and 14.6% (39 out of 268 students) missed 16 or more classes (equivalent in learning time to an entire program of *Discovery*) in a semester (**Fig 2h**.

*Discovery* EE students (**Fig 3**), roughly by definition, obtained lower course grades (p < 0.0001, **Fig 3a**) and higher final *Discovery* grades (p = 0.0004, **Fig 3b**) than non-EE students. This cohort of students exhibited program grades significantly higher than classmates (**Fig 3d-h**) in every category with the exception of essays, where they performed to a significantly lower degree (p = 0.097; **Fig 3c**). There was no statistically significant difference in EE vs. non-EE student classroom attendance (p = 0.85; (**Fig 3i-j**).. There were only 4 single day absences in *Discovery* within the EE subset; however, this difference was not statistically significant (p = 0.074).

**Fig 3.**
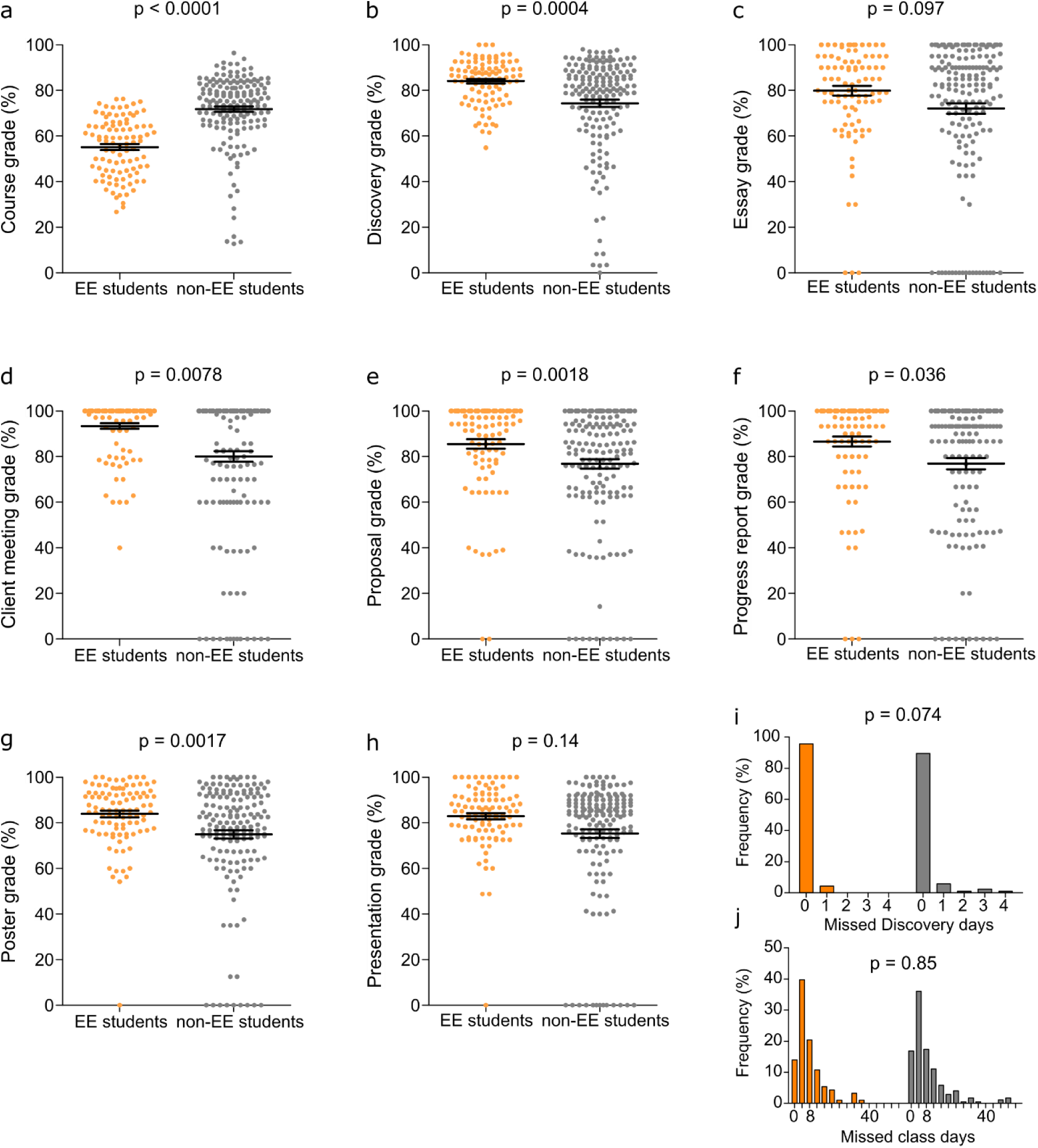
Performance of exceeds expectations student subset. The “Exceeds Expectations” (EE) subset of students (defined as those who received a combined *Discovery* grade ≥ 1 SD (18.0%) higher than their final course grade) performed (a) lower on their final course grade and (b) higher in the *Discovery* program as a whole when compared to their classmates. (d-h) EE students received significantly higher grades on each *Discovery* deliverable than their classmates, except for their (c) introductory essays and (h) final presentations. The EE subset also tended (i) to have a higher relative rate of attendance during *Discovery* sessions but no difference in (j) classroom attendance. N = 99 EE students and 169 non-EE students (268 total). Grade data expressed as mean ± SEM.

*Discovery* MT students (**Fig 4**), although not receiving significantly higher grades in class than students participating in the program only one time (p = 0.29, **Fig 4a**), were observed to obtain higher final *Discovery* grades than single-term students (p = 0.0067, **Fig 4b**). However, MT students only performed significantly better on the progress report (p = 0.0021; **Fig 4f**), with trends of higher performance for their initial proposals and final presentations (p = 0.081 and 0.056, *respectively*; **Fig 4e & 4h**); all other deliverables were not significantly different between MT and non-MT students (**Fig 4c-d & 4g**). Attendance in *Discovery* (p = 0.22) was also not significantly different between MT and non-MT students, although MT students did miss significantly less class time (p = 0.010) (**Fig 4i-j**).

**Fig 4.**
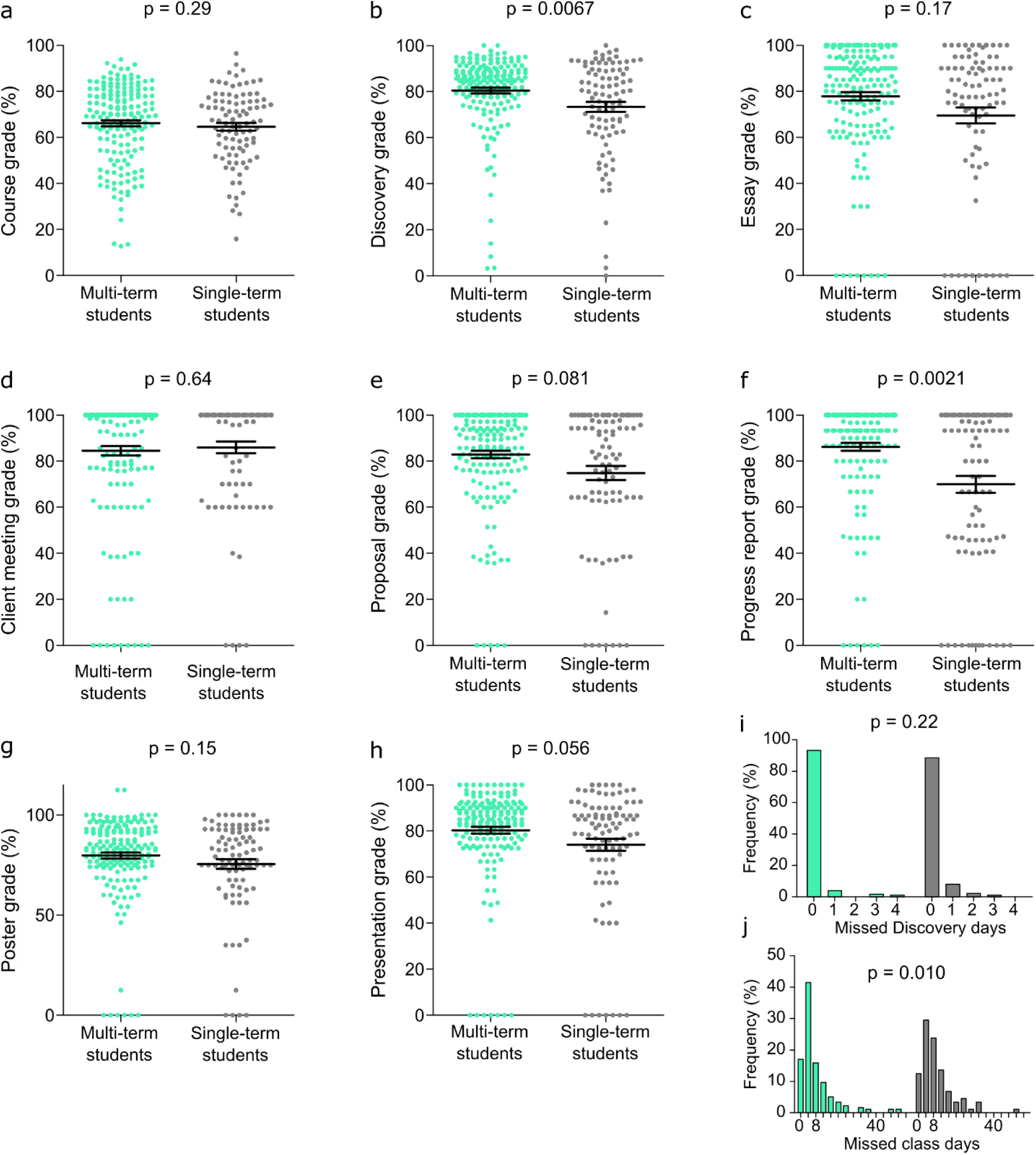
Performance of multi-term student subset. The “multi-term” (MT) subset of students (defined as having attended more than one semester of *Discovery*) demonstrated favourable performance in *Discovery*, (a) showing no difference in course grade compared to single-term students, but (b outperforming them in final *Discovery* grade. Independent of the number of times participating in *Discovery*, MT students did not score significantly differently on their (c) essay, (d) client meeting, or (g) poster. They tended to outperform their single-term classmates on the (e) proposal and (h) final presentation, and scored significantly higher on their (f) progress report. MT students showed no statistical difference in (i) *Discovery* attendance, but did show (j) higher rates of classroom attendance than single-term students. N=174 MT instances of student participation (76 individual students) and 94 single-term students. Grade data expressed as mean ± SEM.

### 2.3 Educator Perceptions

Qualitative observation in the classroom by high school educators emphasized the value students independently placed on program participation and deliverables. Throughout the term, students often prioritized *Discovery* group assignments over other tasks for their STEM courses, regardless of academic weight and/or due date. Comparing within this student population, educators spoke of difficulties with late and incomplete assignments in the regular curriculum but found very few such instances with respect to *Discovery*-associated deliverables. Further, educators speculated on the good behaviour and focus of students in *Discovery* programming in contrast to attentiveness and behaviour issues in their school classrooms. Multiple anecdotal examples were shared of renewed perception of student potential; students that exhibited poor academic performance in the classroom often engaged with high performance in this inquiry-focused atmosphere. Students appeared to take a sense of ownership, excitement and pride in the setting of group projects oriented around scientific inquiry, discovery, and dissemination.

### 2.4 Student Perceptions

Students were asked to consider and rank the academic difficulty (scale of 1-5, with 1 = not challenging and 5 = highly challenging) of the work they conducted within the *Discovery* learning model. Considering individual *Discovery* terms, at least 91% of students felt the curriculum to be sufficiently challenging with a 3/5 or higher ranking (Term 1: 87.5%, Term 2: 93.4%, Term 3: 85%, Term 4: 93.3%, Term 5: 100%), and a minimum of 58% of students indicating a 4/5 or higher ranking (Term 1: 58.3%, Term 2: 70.5%, Term 3: 67.5%, Term 4: 69.1%, Term 5: 86.4%) (**Fig 5a**).

**Fig 5.**
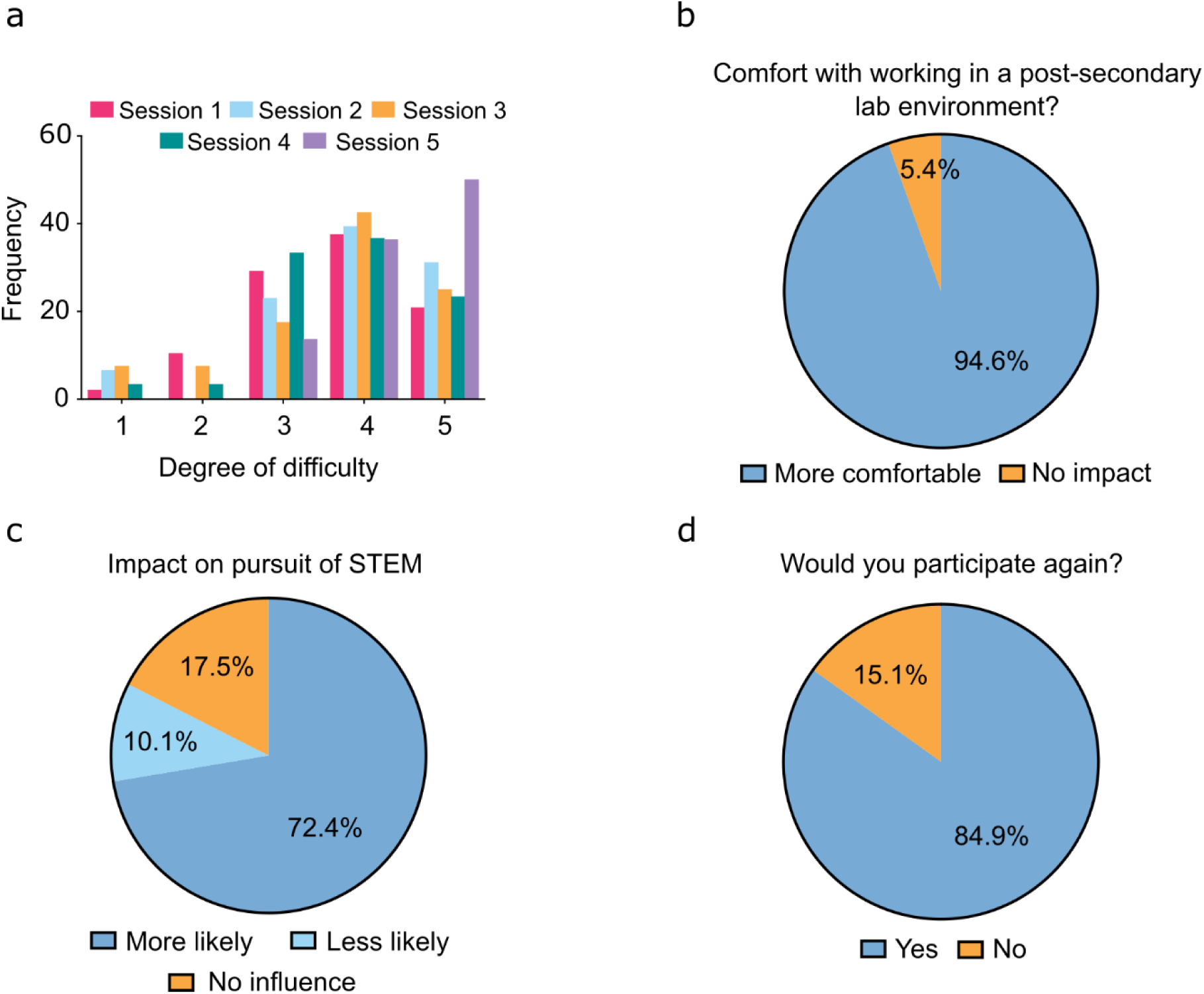
Student survey responses following participation in *Discovery* programming. (a) Histogram of relative frequency of perceived *Discovery* programming academic difficulty ranked from not challenging (1) to highly challenging (5) for each session demonstrated the consistently perceived high degree of difficulty for *Discovery* programming (total responses: 223). (b) Program participation increased student comfort (94.6%) with navigating lab work in a university or college setting (total responses: 220). (c) Considering participation in *Discovery* programming, students indicated their increased (72.4%) or decreased (10.1%) likelihood to pursue future experiences in STEM as a measure of program impact (total responses: 217). (d) Large majority of participating students (84.9%) indicated their interest for future participation in *Discovery* (total responses: 212). Students were given the opportunity to opt out of individual survey questions, partially completed surveys were included in totals.

The majority of students (94.6%) indicated they felt more comfortable with the idea of performing future work in a university STEM laboratory environment given exposure to university teaching facilities throughout the program (**Fig 5b**). Students were also queried whether they were i) more likely, ii) less likely, or iii) not impacted by their experience in the pursuit of STEM in the future. The majority of participants (> 82%) perceived impact on STEM interests, with 72.4% indicating they were more likely to pursue these interests in the future (**Fig 5c**). When surveyed at the end of term, 84.9% of students indicated they would participate in the program again (**Fig 5d**).

## 3 Discussion

We have described an inquiry-based framework for implementing experiential STEM education in a BME setting. Using this model, we engaged 268 participants (170 individual students) over five terms in project-based learning wherein students worked in peer-based teams under the mentorship of U of T trainees to design and execute the scientific method in answering a relevant research question. Collaboration between high school educators and *Discovery* instructors allowed for high school student exposure to cutting edge BME research topics, participation in facilitated inquiry, and acquisition of knowledge through scientific discovery. All assessments were conducted by high school educators and constituted a fraction (10-15%) of the overall course grade, instilling academic value for participating students. As such, students exhibited excitement to learn as well as commitment to their studies in the program.

Through our observations and analysis, we suggest there is value in differential learning environments for students that struggle in a knowledge acquisition-focused classroom setting. In general, we observed a high level of academic performance in *Discovery* programming (**Fig 2a**), which was highlighted exceptionally in EE students who exhibited greater academic performance in *Discovery* deliverables compared to normal coursework (> 18% grade improvement in relevant deliverables). We initially considered whether this was the result of strong students influencing weaker students; however, group organization within each course suggests this is not the case (**Fig 2d**). With the exception of one class in one semester (24 participants assigned by their educator), students were allowed to self-organize into working groups and they chose to work with other students of relatively similar academic performance (as indicated by course grade), a trend observed in other studies^27,28^. Remarkably, EE students not only excelled during *Discovery* when compared to their own performance in class, but this cohort also achieved significantly higher average grades in each of the deliverables throughout the program when compared to the remaining *Discovery* cohort (**Fig 3**). This data demonstrates the value of an inquiry-based learning environment compared to knowledge focused delivery in the classroom in allowing students to excel. It is a well-supported concept that students who struggle in traditional settings tend to demonstrate improved interest and motivation in STEM when given opportunity to interact in a hands-on fashion, which supports our outcomes^3,29^. Furthermore, these outcomes clearly represent variable student learning styles, where some students benefit from a greater exchange of information, knowledge and skills in a cooperative learning environment^30^. The performance of the EE group may not be by itself surprising, as the identification of the subset by definition required high performers in *Discovery* who did not have exceptionally high course grades; in addition, the final *Discovery* grade is dependent on the component assignment grades. However, the discrepancies between EE and non-EE groups attendance suggests that students were engaged by *Discovery* in a way that they were not by regular classroom curriculum.

In addition to quantified engagement in *Discovery* observed in academic performance, we believe remarkable attendance rates are indicative of the value students place in the differential learning structure. Given the differences in number of *Discovery* days and implications of missing one day of regular class compared to this immersive program, we acknowledge it is challenging to directly compare attendance data and therefore approximate this comparison with consideration of learning time equivalence. When combined with other subjective data including student focus, requests to work on *Discovery* during class time, and lack of discipline/behaviour issues, the attendance data importantly suggests that students were especially engaged by the *Discovery* model. Further, we believe the increased commute time to the university campus (students are responsible for independent transit to campus, a much longer endeavour than the normal school commute), early program start time, and students’ lack of familiarity with the location are non-trivial considerations when determining the propensity of students to participate enthusiastically in *Discovery*. We feel this suggests the students place value on this team-focused learning and find it to be more applicable and meaningful to their interests.

Given post-secondary admission requirements for STEM programs, it would be prudent to think that students participating in multiple STEM classes across semesters are the ones with the most inherent interest in post-secondary style STEM programs. The MT subset, representing students who participated in *Discovery* for more than one semester, averaged significantly higher final *Discovery* grades. The increase in the final *Discovery* grade was observed to result from a general confluence of improved performance over multiple deliverables and a continuous effort to improve in a STEM curriculum. This was reflected in longitudinal tracking of *Discovery* performance, where we observed a significant trend of improved performance. Interestingly, the high number of MT students who were included in the EE group suggests that students who had a keen interest in science enrolled in more than one course and in general responded well to the inquiry-based teaching method of *Discovery*, where scientific method was put into action. It stands to reason that even if they do not perform well in their specific course, students interested in science will continue to take STEM courses and will respond favourably to opportunities to put classroom theory to practical application.

The true value of an inquiry-based program such as *Discovery* may not be based in inspiring students to perform at a higher standard in STEM within the high school setting, as skills in critical thinking do not necessarily translate to knowledge-based assessment. Notably, students found the programming equally challenging throughout each of the sequential sessions, perhaps somewhat surprising considering the increasing number of repeat attendees in successive sessions (**Fig 5a**). Regardless of sub-discipline, there was an emphasis of perceived value demonstrated through student surveys where we observed indicated interest in STEM and comfort with laboratory work environments, and desire to engage in future iterations given the opportunity. Although non-quantitative, we perceive this as an indicator of significant student engagement, even though some participants did not yield academic success in the program and found it highly challenging given its ambiguity. Further, we observed that students become more certain of their direction in STEM, correlating with preliminary trends of increased post-secondary application rates by *Discovery* graduates (data not shown); further longitudinal study is warranted to make claim of this result. At this point in our assessment we cannot effectively assess the practical outcomes of participation, understanding that the immediate effects observed are subject to a number of factors associated with performance in the high school learning environment. Future studies that track graduates from this program will be prudent, in conjunction with an ever-growing dataset of assessment, to continue to understand the expected benefits of this inquiry-focused and partnered approach. Altogether, a multifaceted assessment of our early outcomes suggests significant value of an immersive and iterative interaction with STEM as part of the high school experience. A well-defined divergence from knowledge-based learning, focused on engagement in critical thinking development framed in the cutting-edge of STEM, may be an important step to broadening student perspectives.

As we consider *Discovery* in a bigger picture context, expansion and implementation of this model is translatable. Execution of the scientific method is an important aspect of citizen science, as the concepts of critical thing become ever-more important in a landscape of changing technological landscapes. Giving students critical thinking and problem-solving skills in their primary and secondary education provides value in the context of any career path. Further, we feel that this model is scalable across disciplines, STEM or otherwise, as a means of building the tools of inquiry. We have observed here the value of differential inclusive student engagement and critical thinking through an inquiry-focused model for a subset of students, but further to this an engagement, interest and excitement across the body of student participants. As we educate the leaders of tomorrow, we suggest use of an inquiry-focused model such as *Discovery* could facilitate growth of a data-driven critical thinking framework.

## 4 Methods

### 4.1 Experimental Design

All students in university-stream Grade 11 or 12 biology, chemistry, or physics at the participating school were recruited into mandatory offerings of *Discovery* over five consecutive terms. Student grade and survey responses were collected pending parent or guardian permission. Educators replaced each student name with a unique coded identifier to preserve anonymity but enable individual student tracking over multiple terms. All data collected was analyzed without any exclusions save for missing survey responses; no power analysis was performed prior to data collection.

### 4.2 Ethics statement

This study was approved by the University of Toronto Health Sciences Research Ethics Board (Protocol # 34825) and the Toronto District School Board External Research Review Committee (Protocol # 2017-2018-20). Acquisition of student data (both post-hoc academic data and survey administration) followed written informed consent of data collection from parents or guardians of participating students. Data was anonymized by high school educators for maintenance of academic confidentiality.

### 4.3 Program overview

In facilitation of *Discovery*, a selected global health research topic was sub-divided into subject-specific research questions (i.e., Biology, Chemistry, Physics) that students worked to address, both on-campus and in-class, during a term-long project. The *Discovery* framework therefore provides students the experience of an engineering capstone design project, and includes a motivating scientific problem (i.e., global topic), a discipline-specific research question, and systematic determination of a professional recommendation addressing the needs of the presented problem.

#### 4.3.1 High school partner

The *Discovery* program evolved to the current model over a two-year period of working with one high school selected from the local public school board. This partner school consistently scores highly (top decile) in the board’s Learning Opportunities Index (LOI). The LOI ranks each school based on measures of external challenges affecting student success, therefore schools with the greatest level of external challenges receive a higher ranking^31^. Consequently, participating students are identified as having a significant number of external challenges that may affect their academic success. In addition, the selected school partner is located within a reasonable geographical radius of our campus (i.e., ∼ 40 min transit time from school to campus). This is relevant as participating students are required to independently commute to campus for *Discovery* hands-on experiences.

#### 4.3.2 Student recruitment

In agreement with school administration and science educators, *Discovery* was incorporated as a mandatory component of course curriculum for senior students (Grade 11 and 12) in university stream Chemistry, Physics, and Biology courses. Students therefore participated as class cohorts to address questions specific to their course discipline knowledge base, but relating to the defined global health research topic (**Fig 1**). At the discretion of each STEM teacher, assessment of program deliverables was collectively assigned as 10-15% of the final course grade for each subject. All students were required to participate; however, students were given opportunity to opt out the research study aspect of this program and parent/guardian consent was required for student data to be collected and collated for research purposes.

#### 4.3.3 Instructional framework

Each program term of *Discovery* corresponds with a five-month high school semester. U of T trainees (*Discovery* instructors) and high school educators worked collaboratively to define a global healthcare theme in advance of each semester. In addition, specific cutting-edge and curriculum-relevant research questions were developed for each discipline to align within both the overall theme and the educational outcomes set by the provincial curriculum ^32^. *Discovery* instructors were consequently responsible for developing and introducing relevant STEM skills, as well as mentoring high school students, for the duration of their projects; high school educators were responsible for academic assessment of all program deliverables throughout the term (**Fig 1**).

During the course of a term, students engaged within the university facilities four times. The first three sessions included hands-on lab sessions while the fourth visit included a culminating symposium for students to present their scientific findings (**Fig 1**). Project execution was supported by U of T trainees who acted as engineering “clients” to mentor student groups in developing and improving their assessment protocols, as well as generating final recommendations to the original overarching questions. On average, there were 4-5 groups of students per discipline (3-4 students per group; ∼20 students/class). *Discovery* instructors worked exclusively with 1-2 groups each term in the capacity of mentor to monitor and guide student progress.

After introducing the selected global research topic in class, educators led students in completion of background research essays. Students subsequently engaged in a discipline-relevant skill-building protocol during their first visit to university teaching laboratory facilities, allowing opportunity to understand analysis techniques and equipment relevant for their assessment projects. At completion of this session, student groups were presented with a discipline-specific research question as well as the relevant laboratory inventory available for use during their projects. Armed with this information, student groups continued to work in their classroom setting to develop group-specific experimental plans. Educators and *Discovery* instructors provided written and oral feedback, *respectively*, allowing students an opportunity to revise their plans in class prior to on-campus experimental execution. Once at the relevant laboratory environment, students executed their protocols in an effort to collect experimental data. Data analysis was performed in the classroom and students learned by trial & error to optimize their protocols before returning to the university lab for a second opportunity for data collection. All methods and data were re-analyzed in class in order for students to create a scientific poster for the purpose of study/experience dissemination. During a final visit to campus, all groups presented their findings at a research symposium, allowing students to verbally defend their process, analyses, interpretations, and design recommendations to a diverse audience including peers, STEM educators, undergraduate and graduate university students, postdoctoral fellows and University of Toronto faculty.

#### 4.3.4 Data collection

Educators evaluated students within their classes on the following associated deliverables: i) global theme background research essay; ii) experimental plan; iii) progress report; iv) final poster content and presentation; and v) attendance. For research purposes, these grades were examined individually and also as a collective *Discovery* program grade. For students consenting to participation in the research study, all *Discovery* grades were anonymized by the classroom educator before being shared with study authors. Each student was assigned a code (known only to the classroom educator) for direct comparison of deliverable outcomes and survey responses.

Survey instruments were used to gain insight into student perceptions of STEM and post-secondary study, as well as *Discovery* program experience and impact (**S2 Appendix II**). High school educators administered surveys in the classroom only to students supported by parental permission. Pre-program surveys were completed at minimum one week prior to program initiation each term and exit surveys were completed at maximum two weeks post-*Discovery* term completion.

### 4.4 Identification and comparison of population subsets

From initial analysis, we identified two student subpopulations of particular interest: students who performed ≥ 1 SD [18.0%] or greater in the *Discovery* portion of the course compared to their final course grade (“EE”), and students who participated in *Discovery* more than once (“MT”). These groups were compared individually against the rest of the respective *Discovery* population (“non-EE” and “non-MT”, *respectively*). Additionally, MT students who participated in three or four (the maximum observed) semesters of *Discovery* were assessed for longitudinal changes to performance in their course and *Discovery* grades. Comparisons were made for all *Discovery* deliverables (introductory essay, client meeting, proposal, progress report, poster, and presentation), final *Discovery* grade, final course grade, *Discovery* attendance, and overall class attendance.

### 4.5 Statistical analysis

Student course grades were analyzed in all instances without the *Discovery* component contribution (ranging from 10% to 15% of final mark depending on class and year) to prevent correlation. Student course grade vs. matched *Discovery* grade was first compared by paired t-test. Student performance (N=268 total students, comprising 170 individuals) in *Discovery* was initially assessed in a linear regression of *Discovery* grade vs. final course grade. Trends in course and Discovery performance over time in students participating 3 or 4 semesters (N=16 and 3 individuals, *respectively*) were also assessed by linear regression. For subpopulation analysis (EE and MT, N=99 instances from 81 individuals and 174 instances from 76 individuals, *respectively*), each data set was tested for normality using the D’Agostino and Pearson omnibus normality test. All subgroup comparisons vs. the remaining population were performed by Mann-Whitney U-test. Data are plotted as individual points with mean ± SEM overlaid (grades), or in histogram bins of 1 and 4 days, *respectively*, for *Discovery* and class attendance. Significance was set at α ≤ 0.05.

## 6 Acknowledgements

The authors would like to acknowledge the support of the many post secondary trainee volunteers from the University of Toronto for their countless hours in program planning and execution. In particular, members of the executive organizing team that have assisted study authors with curriculum development to date include Genevieve Conant, Sherif Ramadan, Daniel Smieja, Rami Saab, Andrew Effat, Serena Mandla, Cindy Bui, Janice Wong, Dawn Bannerman, Allison Clement, Shouka Parvin Nejad, Nicolas Ivanov, Jose Cardenas, Huntley Chang, Romario Regeenes, Dr. Henrik Persson, Ali Mojdeh, Nhien Tran-Nguyen, Ileana Co, and Jonathan Rubianto. We further acknowledge the staff of the science department at George Harvey Collegiate Institute for collaboration with program development, execution and administrative support. In particular, Jennifer Alves and Figen Irumekhai (staff) and principals Anthony Vandyke and Sam Micelli (administration) have assisted study authors with programming ideation, delivery, and data collection. *Discovery* has grown with the continued support of Dr. Christopher Yip (Dean, Faculty of Applied Science and Engineering; former IBBME Director), with the support of IBBME Directors (Dr. Craig Simmons and Dr. Warren Chan), and logistical assistance of the IBBME Undergraduate Programs Office administrator (Brittany Lawrence, Brittany Lauton, Andrew Novoselac, Carina Kamango Esmael and Ivy Hon). *Discovery* facility support has been provided by Dr. Max Giuliani, Dr. Lindsey Fiddes, Nguyen Hoang, Alexander Dean, Lily Jeon, Tommy Pham, and undergraduate work study students Michael Belhu, Rafia Kouser and Shreyashi Saha. We also thank Benjamin Rocheleau and Madeleine Rocheleau for contributions to data collation and Payal C. Patel for advisement on structural program improvements. This program is financially supported by IBBME and the National Science and Engineering Research Council (NSERC) PromoScience program as part of the IBBME “Igniting Youth Curiosity in STEM” initiative (PROSC 515876-2017) co-directed by DMK and Dr. Penney Gilbert. LDH and NIC were supported by Vanier Canada graduate scholarships from the Canadian Institutes of Health Research and NSERC, *respectively*. DMK holds a Dean’s Emerging Innovation in Teaching Professorship in the Faculty of Engineering & Applied Science at the University of Toronto.

## 8 Supplementary Materials

S1 Appendix I: Sample teaching materials for one representative term of *Discovery*

S2 Appendix II: Entrance and exit student surveys for program assessment of *Discovery*

S3 Appendix III: Mark breakdown for student assessment in *Discovery* programming

