## Appendix 1 for "Enhancing senior high school student engagement and academic performance using an inclusive and scalable inquiry-based program"

### S1 Appendix: Sample teaching materials for one term of IBBME Discovery

### Introduction

The goal of IBBME is to provide students with a real challenge in biomedical engineering, for which an introductory skill lab provides them with the technical ability to address the challenge in some way. In addition to the subject-specific technical skills that students will gain, the curriculum is also designed to give students highly transferable general scientific and engineering skills including:

- Literature review
- Experimental design
- Prototyping
- Data collection
- Validation
- Interpretation

For each subject (i.e. biology, chemistry, and physics), we have included:

- An outline of the research project for the purposes of logistical and curricular planning, and indicating the nature of the challenge, research question, expected methodology, and learning outcomes for students
- A skill lab protocol in the style of a standard 3-hour undergraduate laboratory, adjusted for technical and theoretical level and designed to introduce experimental and analytical techniques necessary to complete the conceived project
- A request for proposal (RFP) presented to the students outlining the motivation underlying the biomedical engineering challenge that the students are expected to contribute to, and to which the students must respond with a formal proposal

At the beginning of the skill lab, students were required by the facility to undergo a brief safety training that covered considerations of the facility in general, as well as safety procedures specific to the techniques and equipment in use and standard emergency procedures. Material safety data sheets (MSDS) for chemicals used were made available to all participants before the day of the skill lab. All graduate instructors participated in all department-mandated safety training modules.

After safety training, students typically took part in a 30 to 60 minute interactive lecture pertaining to the grand challenge of the semester, as well as introducing the theory and technical knowledge required to complete the skill lab protocol for the rest of the day. Students had been provided the skill lab protocol before attending, and they were given the rest of the day to complete the protocol as written, with instructor assistance as needed. At the end of the day, the RFP outlining the challenge was distributed to the students, and time was taken to discuss with students the nature of the project and potential avenues of inquiry.

Here we have provided sample documents for a past semester of Discovery, including outlines, skill labs, and RFPs for all three subjects. We have also included the template students use to complete the proposed research plan. The central theme of the semester was “Cardiovascular Disease: Engineering diagnostics and treatments”. This served to constrain project directions, rationalize the skills and techniques covered, and motivate students.

### Biology

#### Biology project outline

**Motivation:** Currently in Canada there are over 1.6 million people with cardiovascular disease, and cardiac health is one of the leading causes of death in the developed world. These patients suffer from a significant reduction in their quality of life as reduction in heart function makes everyday life difficult. Although healthcare continues to advance treatment, there is a need for continued focus on preventing, identifying and treating this prominent health problem.

Cardiotoxicity effects are the leading cause of pharmaceutical drug withdrawal from markets; and delays in regulatory approval. Thus, cardiac safety assessment has become an important regulatory step in drug development. Inappropriate drug formulations or excessive drug exposure can lead to cardiac toxic effects that include arrhythmia, cardiac electrical instability, and sudden cardiac death. Some of these toxicity effects can be regulated in a time-dependent and dose-dependent manner. We want to investigate the cardiotoxicity effects of an unknown chemical compound, Substance X; however, we will require the expertise of cell biologists to conduct experiments comparing Substance X to known cardiac toxic drugs.

**Research question:** How can we characterize the toxic effects of chemical compounds on cardiac cells?

**Rationale:** Knowledge of cardiac cell characteristics (beating frequency, morphology, calcium oscillations, etc.) allows us to distinguish between cardiac cells that are functioning normally and cardiac cells that are beginning to fail. Once there is an understanding of the differences between healthy and dysfunctional cardiac cells, we can start to evaluate the potential cardiac toxicity effects of chemical compounds on cardiac cells.

**Skill-based activity:** A full day of introductory programming will include:

- Laboratory Biosafety (lecture, demonstration, quiz)

- Introduction to Pipetting & Serial Dilutions (lecture, pipetting practice)
- Cell Morphology and Microscopy (lecture, microscopy demo, cell imaging)
- Cell Staining - live/dead, DAPI (lecture, cell imaging)

**Required equipment:** Students will be given plates of cardiac cells, access to fluorescence microscopes, and a chemical inventory to characterize the cardiac tissue and explore how cell characteristics are impacted by various chemical compounds.

**Expected outcomes:** By the end of the module, students will be able to:

- Become familiar with basic laboratory practices and techniques
- Understand cell properties and how to measure them
- Develop testable hypotheses and experiments to prove or disprove them
- Perform fluorescence microscopy and imaging

**Relevant literature topics:** Further reading that may provide a deeper understanding of material covered in this research project can be done by searching relevant topics such as:

- How to use a pipette and perform serial dilutions
- Cell structures and properties of (cardiac) cells
- Changes to cardiac cells at the onset of cardiac cell dysfunction
- Common cellular dyes and how to prepare them for use in cell cultures
- How a fluorescence microscope works
- Components and format for writing a scientific report

#### Biology skill lab

Questions to consider during lecture:

*Why do we culture and experiment on cells?*

*What are common good practices in sterile technique?*

*What is the purpose of cell culture medium?*

*What is fluorescence microscopy used for? What does it mean to "label" something for fluorescent microscopy?*

*What is phase-contrast microscopy used for?*

*How do cells gain and use energy? Where does the energy source come from?*

*At what temperature do we culture mammalian cells? Why else do we keep them in an incubator?*

*What is the aim of sterile technique? Why do we use it?*

*What is experimental variability? Why do we take replicate measurements in an experiment?*

*How large are most mammalian cells? What happens to their apparent size as we increase magnification on a microscope?*

Sterile technique and pipetting practice

<u>Materials</u>

Ultrapure water
5 mmol/L blue dye stock
Microplates
Pipettes and pipette tips
Epitubes

1. While working in the Class II Biosafety Cabinet, and with sterile technique as demonstrated by your instructors, serially dilute the 5 mmol/L blue dye stock solution.

2. Set up 8 Epitubes and label them # 1-8.

3. Pipette 100 µL of the stock solution into a tube # 1 with 900 µL of water by depressing to the second stop, and gently mix by pipetting up and down above the first stop (be careful not to draw solution into the barrel of the pipette!). What concentration do you have now? Label the Epitube with the new concentration.

4. Transfer 100 µL from tube # 1 to tube # 2. Add 900 µL of water to tube # 2. Mix well as before. What concentration do you have now? Label the Epitube with the new concentration of dye solution.

5. Repeat this serial dilution with tubes # 3-8.

*What is the concentration of dye within each tube? What happens to the colour of each solution?*

Protocol: Cell culture and microscopy practice

For today's lab, you will be working with HeLa cells. You aren't expected to know the answers to the questions in the protocol ahead of time. Feel free to discuss with your instructors, and make sure to ask any questions you have. There are no stupid questions in a biology lab!

*What do you know about these cells? Where do they come from?*

**Cell Culture**

1. Obtain two dishes of cells from your instructor. Label them both with your group number or names, and mark one as “control” and one as “room temperature”.
2. Replace the cell culture media in both dishes as directed by your instructor, using sterile technique. Return the “control” plate to the incubator and leave the “room temperature” plate in the sterile flow hood before you go to lunch.

*Based on what you know about cell culture, what do you think will happen to these cells? How could you test this hypothesis?*

3. When you return from lunch, obtain the stock solution tubes of ethidium homodimer (EthD) and calcein-AM from your instructor. Make sure to leave the aluminum foil on the tubes.

*Why is it important to shield these solutions from light?*

4. Using sterile technique, add 5 µL of EthD and 2.5 µL of calcein-AM to 5 mL of phosphate-buffered saline (PBS) in a new 15 mL Falcon tube. This is your dye solution.
5. Remove the cell culture medium from both plates and add 2.5 mL of your dye solution to both plates. Allow them to incubate for 30-45 min.

**Fluorescent Microscopy**

1. With the instructor’s help, look at your cells using the microscope. Image your plates using brightfield, phase contrast, and red and green fluorescent channels. Record representative images (at least 3 replicates per plate) in all modes using both 10X and 40X objectives. Note that you need to capture all channels in the same place before moving your sample.

2. For each filter and magnification, use the same exposure time (i.e. all 10x TX2 images should have the same exposure time. This can be different from the 40x TX2 exposure time and from the 10x L5 exposure time).

3. In total, you should take 24 images (control: 3 brightfield, 3 phase contrast, 3 green, 3 red; room temp: 3 brightfield, 3 phase contrast, 3 green, 3 red).

4. When saving your image, save it as a tif file and make sure the filename includes the following details: Cell name, treatment, objective magnification, filter and exposure time. For instance: *HeLa_room temp_10x, L5, 80 ms.tif*

5. Save the images in a folder with your name or group number and today's date.

**TIP:** do *not* include a scale bar in the exported image. You can add a scale to the image in ImageJ without hard-coding a scale bar into the raw image.

*What is the purpose of each channel with which you are taking images?*

*Why is it important to capture images in all channels in one area?*

*Why is it important to keep exposure time constant between images with same filter and magnification? What would happen otherwise?*

**<u>Densitometry using ImageJ</u>**

1. For specific details and guides, see the "ImageJ" section below.

2. Open ImageJ on your computer.

3. Open an image by dragging & dropping it on the ImageJ interface. Before starting to work with your images, it is important to set the scale of the image, i.e. tell the software how large it is in real world units. To do this, click Analyze>Set scale and enter the appropriate information from the table in Appendix 2.

4. Cell counting: From your 10X images, use a particle counting tool (Plugins > Analyze > Cell Counter) to count how many cells are fluorescing red, and how many are fluorescing green. Calculate the percentage of dead cells for each sample.

5. Make a composite image incorporating your phase contrast, red, and green channels (Image > Color > Merge Channels). Add a scale bar based on the scale information you added in step 8: Analyze>Tools>Scale Bar.

6. Intensity measurement**:** Set your image measurements: Analyze > Set Measurements, enable “Area”, “Integrated density” and “Display Label”. (The label is not necessary but helps you keep track of which image the data comes from). For a 40X green fluorescent image, select the interior of a single cell by tracing its perimeter with the Magic Wand and/or Freehand Selection tool(s). Take a measurement of the cell (M or Analyze > Measure). This will produce both an area and density measurement and display them in a Results table. Repeat for ten cells in each treatment. Save your Results.

7. Analysis: Use excel to open your results file. Divide each cell’s RawIntDen value by its area. List these intensity/area ratios by each treatment.

8. Calculate the mean (Excel: “=AVERAGE([cell1]:[cell10])”) and standard deviation (Excel: “=STDEV([cell1]:[cell10])”) of both treatments.

9. Run a two-sample t-test (Excel: “=TTEST([cell1a]:[cell10a],[cell1b]:[cell10b],2)”) to see if your treatments are significantly different.

*What is the purpose of density? Area? Density over area (normalized density)?*

*What does statistical significance mean? How is it determined?*

*Would you expect your two treatments to differ in normalized density? Why or why not? What other parameters might you expect to be different?*

**ImageJ**

ImageJ is a free to use, open source image manipulation and analysis software which is very popular in the microscopy community. It has been around since 1997 and its open source status has enabled researchers to create their own plugins, expanding the usefulness of the program. If you want to try it at home, you can find installation instructions here: https://imagej.nih.gov/ij/download.html.

**Set scale:** The scale of an image is a product of camera pixel size and the microscope's total magnification and is an important aspect of microscopy; it is required to know how large anything in your image actually is. To encode an image's scale in the data for the image click Analyze>Set Scale and fill out the dialog box according to this table:

| **Obj.** | **Distance in pixels** | **Known distance** | **Pixel aspect ratio** | **Unit** |
|---|---|---|---|---|
| 5x | 1 | 0.41 | 1 | um |
| 10x | 1 | 0.20 | 1 | um |
| 20x | 1 | 0.10 | 1 | um |
| 40x | 1 | 0.05 | 1 | um |

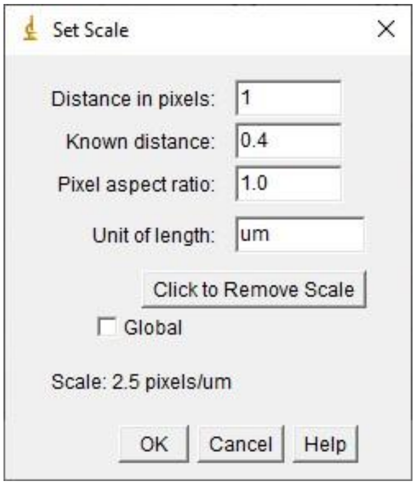

*Fig. 1: Set scale dialog and validated scale distances for microscope objectives in ImageJ.*

**Cell counting:** Use the plugin “Cell counter” to count cells in your images. Open the plugin: Plugins>Analyze>Cell counter and then click on your image, making it active. Click initialize in the plugin window, select a counter type and make sure you have one of the selection tools active (e.g. click on the “freehand selection” tool in the ImageJ toolbar). Click on all cells of interest.

If you make a mistake, you can press “delete” in the plugin and the last click of that type will be removed. If you check the box “delete mode”, you can click in the image to delete the closest marker of that type.

**Obsolete:** In newer installations of ImageJ, the Cell Counter plugin has been removed and is replaced by the native Multipoint Tool which is even easier to use. The plugin should be available on all lab computers but if you install ImageJ at home, this might not be the case. To run the multipoint tool, simply launch it from the toolbar, select a colour and cell type number and click away. (Alternately, try to open the plugin as normal and ImageJ will automatically launch Multipoint tool for you.)

**Tracing cells:**

**Wand tool:** Using the Wand Tool, you can click on a pixel and all connected pixels of similar intensity will be selected. This is good if there is high contrast between cells (e.g. a darker area between). If you double click the wand tool, you can try different methods to determine “neighbouring” pixels and you can set tolerance for variations. A tolerance of 0 means pixels need to have the same intensity to be selected. Increasing the tolerance gives more leeway and pixels of similar intensity will be selected. This can be done live by selecting a cell and moving the slider.

**Freehand selection:** if the image is such that the wand tool is working poorly, cells can be manually traced with the freehand selection tool. Select the tool, click and hold the mouse button while tracing the outline of your cell.

**Modifications:** if you make a bad drawing, rather than starting over from scratch, you can modify it. By holding Shift while tracing, the new area will be added to the original area. (The two areas do not need to overlap.) By instead holding Alt while tracing, you can subtract areas.

These modifications can be combined with the wand tool: select a cell using the wand tool and add or subtract areas with the freehand tool.

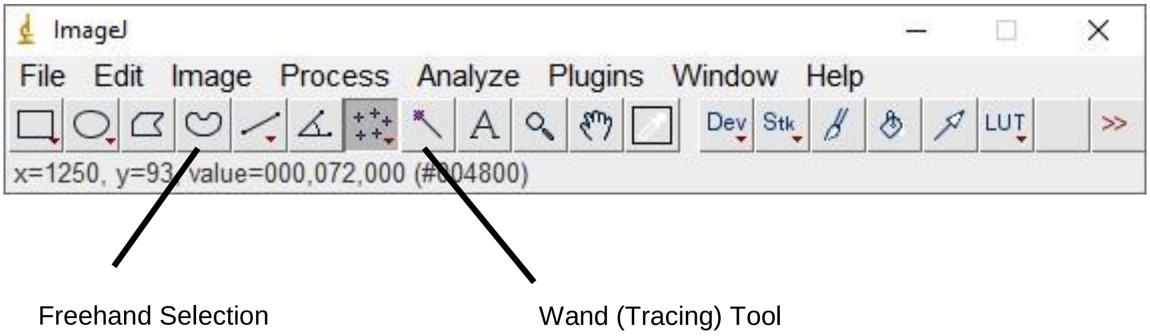

*Fig. 2: Selection tools in ImageJ’s simple interface.*

#### Biology request for proposal

Currently in Canada there are over 1.6 million people with cardiovascular disease, and cardiac health is one of the leading causes of death in the developed world. These patients suffer from a significant reduction in their quality of life as reduction in heart function can make everyday life difficult. Although healthcare continues to advance treatment, there is a need for continued focus on preventing, identifying and treating this prominent health problem. We are looking for the next generation of great minds to help us improve the lives of these people around the world!

Cardiotoxicity effects are the leading cause of pharmaceutical drug withdrawal from markets and delays in regulatory approval. Thus, cardiac safety assessment has become an important regulatory step in drug development. Inappropriate drug formulations or excessive drug exposure can lead to toxic effects including arrhythmia, cardiac electrical instability, and sudden cardiac death. Some of these toxicity effects can be regulated in a time-dependent and dose-dependent manner. **We want to investigate the cardiotoxic effects of a newly developed chemical compound, Substance X. However, we will require the expertise of cell biologists to conduct experiments comparing Substance X to known chemicals and/or drugs.**

We will be collecting proposals from multiple teams and will consider all options at a collective symposium. Please submit your project proposal using the RFP template provided. We have attached a list of the materials available in our facilities, so please take a look at these as you build your experimental plan. These will be reviewed by IBBME representatives who will meet with you to finalize the experimental details. You will then utilize our facilities to carry out your project and deliver a final presentation in poster format (template provided). We are looking forward to seeing the innovative solutions you will develop!

**Notes:**

**Be aware of experimental constraints**

*Time:* Plan for 5h in the laboratory per visit (2 visits total) to do your experiments and as well as some data analysis

*Materials & Equipment:* See attached inventory list

Proposals requiring materials that are not on the inventory list may also be approved on a case-by-case basis

**Do your research**

- How do we define the best dose for a drug?
  {e.g. dose concentration, number of doses, time between doses, etc.}
- What are properties used to characterize a drug's cardiotoxic effects?
  {e.g. effects on cell health/viability, cell morphology, cell behaviour, etc.}
- What techniques exist to measure these properties?
  {e.g. changes in cell adherence, changes in cell morphology/phenotype, changes in cell viability staining results, etc.}

**Choose a cellular property to investigate as well as a set of known chemicals and/or drugs. Describe the procedures you would perform in order to characterize the potentially cardiotoxic effects of Substance X relative to known substances.**

- Provide justification for your decisions
- Keep your proposal simple, you are limited in time and other resources

**If your proposal is approved, be prepared to make it happen!**

- Consider developing a spreadsheet/table for recording your observations

- Contact lenses are not allowed in the laboratory (please wear glasses)

Long pants, closed-toed shoes, and socks are required to work in the laboratory

### Chemistry

#### Chemistry project outline

**Motivation:** Currently in Canada there are over 1.6 million people with cardiovascular disease, and cardiac health is one of the leading causes of death in the developed world. These patients suffer from a significant reduction in their quality of life as reduction in heart function makes everyday life difficult. Although healthcare continues to advance treatment, there is a need for continued focus on preventing, identifying and treating this prominent health problem.

Metoprolol is a beta-blocker that is frequently used to treat high blood pressure and tachycardia (chronically elevated heart rate). It can be taken by pill or intravenously but is a “dirty drug” in that it has numerous side effects, including depression, insomnia, fatigue, dangerously low heart rate, or respiratory depression, among others. Part of this toxicity can be avoided by tightly regulating the concentration of the drug within the body. We are working on bringing a new metaprolol treatment to market in either pill or implant form but need chemical engineering expertise to optimize the release of the drug within safe limits.

**Research question:** Can a hydrogel’s properties be engineered to release a drug within safe limits over time?

**Rationale:** Knowledge of cardiac cell characteristics (beating frequency, morphology, calcium oscillations, etc.) allows us to distinguish between cardiac cells that are functioning normally and cardiac cells that are beginning to fail. Once there is an understanding of the differences between healthy and dysfunctional cardiac cells, we can start to evaluate the potential cardiac toxicity effects of chemical compounds on cardiac cells.

**Skill-based activity:** Students will compare prepared gelatin to alginate beads in their release kinetics of food colouring using a spectrophotometer and standard curve. Knowledge of the Beer-Lambert law and chemical equilibria will be necessary. Students will gain experience with all relevant techniques and will learn to use Excel graphing to draw quantitative conclusions. Students will use FCF Brilliant Blue #1 (blue food colouring) for all activities. First, they will be required to create a serial dilution series of stock dye to find a dilution suitable for reading in a spectrophotometer. By using a known extinction coefficient, they will be able to calculate the stock concentration. Next, they will need to create a 5-point linear standard curve within the optimal range of the spectrophotometer based on their understanding of the relationship between absorbance and concentration. Finally, they will use this curve to measure the release of Blue #1 from premade alginate beads into media over time.

**Project:** Students will create hydrogels from alginate and/or gelatin and measure their release of blue dye over time. They will be able to modulate gel composition and density, form (size/shape), and initial amount of dye loaded. Using a standard curve and timepoint measurements, they will generate release profiles that will be loaded into a premade mathematical model of metoprolol absorption and clearance from the human body. This will allow them to assess the efficacy and safety of their hydrogel drug delivery system.

Required equipment

- Gelatin
- Alginate
- Calcium chloride
- Microplates
- Pipette tips
- Beakers

- Food colouring (Blue 1: 628 nm, 1.30*10^5 M-1cm-1, 792.844 g/mol; Yellow 5: 425 nm, 2.73*10^4 M-1cm-1, 534.36 g/mol; Red 40: 504 nm, 2.59*10^4 M-1cm-1, 496.42 g/mol)
- Miscellaneous materials for casting shapes (glass slides, tape, PDMS strips, etc)
- Pipettes
- Spectrophotometer

**Expected outcomes:** At the end of this module, students will be able to:

- Apply concepts and use common equations in equilibrium and the Beer-Lambert Law.
- Understand the theory and limitations of a spectrophotometer.
- Understand hydrogel structure and function.
- Design a proposal to tailor the rate of drug release within a specified range, based on modifying common material properties of a hydrogel (size, shape, polymer content, starting concentration).

**Relevant literature topics:** Custom protocols and presentations will be provided. Readings on equilibrium, conservation of matter, etc. in their own textbooks will be helpful for review.

#### Chemistry skill lab

To be completed during lecture:

*Metoprolol is a beta-blocker. What is the role of a beta-blocker in cardiac medicine?*

*What are three of the side-effects of metoprolol?*

*In general, how can we avoid side-effects while gaining desirable effects of a drug?*

*What properties of a hydrogel can be changed to tune drug release?*

Today's activity:

Today we will be modeling drug concentrations and controlled release using a blue dye (FCF Brilliant Blue #1, or blue food colouring). Blue #1 shows peak absorbance at 628 nm, has a molar mass of 792.84 g/mol, and an extinction coefficient of 130 000 L/mol/cm.

**You should be writing down all steps and data obtained in your notebook. A volunteer must sign off on this protocol/worksheet and your notebook before you leave for the day. All of the techniques and calculations you do today will be important for your design project.**

*Sketch a plot of absorbance vs. wavelength for Blue #1:*

*Why should we measure absorbance at the peak wavelength?*

1. Place 10x 1.6 mL epitubes in a rack. Add 2 drops of blue food colouring to one. This is your working dye stock solution.
2. To another 1.6 mL epitube, pipette 10 µL of blue food colouring using a P10 or P20 pipette. Add 90 µL of ultrapure water.

*What is the dilution of this solution in relation to the original dye stock?*

*Why should we use ultrapure water for spectrophotometry?*

3. Add 10 µL of the solution made in (2) to 90 µL of ultrapure water in a new tube.

*What is the dilution of this solution in relation to the original dye stock?*

4. Add 50 µL of the solution made in (3) to 450 µL of ultrapure water in a new tube.

*What is the dilution of this solution in relation to the original dye stock?*

5. Repeat steps 2-4 two more times until you have 3 **triplicate** samples. Pipette 200 µL of this solution each into wells A1, A2, and A3 of a spectrophotometry microplate
6. With the help of a volunteer, measure the absorbance at 628 nm of these wells.

**Record your three absorbances here as well as in your lab notebook**

7. Using $A = \varepsilon \times C \times l$, compute the current (diluted) concentration of Blue #1 in each of the three triplicate samples you measured. Multiply by the dilution factor to give three replicate values for the initial stock concentration of FCF Brilliant Blue #1.

**Record the three absorbances here, as well as their mean (average) and standard deviation $SD = \sqrt{(\frac{(repl_1 - mean)^2 + (repl_2 - mean)^2 + (repl_3 - mean)^2}{2})}$. Record all computations in your lab notebook.**

*What is the purpose of a standard deviation?*

8. Make another 1 in 1000 dilution by diluting 10 µL of the original dye stock in 90 µL water, 20 µL of this dilution in 180 µL water, and 120 µL of this dilution in 1080 µL water (2x 540 µL). Repeat this process once to generate triplicates. Be careful to not mix up tubes in your rack.

*Why should we dilute samples stepwise, and not by a 1 in 1000 factor in a single step?*

9. Make a serial dilution series for a standard curve by diluting your triplicate 1 in 1000 stocks. From the tube made in (8), add 960 µL to 240 µL water.
10. Add 960 µL of the solution made in (9) to 240 µL water. Repeat this serial dilution process two more times in this dilution series, and then produce two more series using the triplicates you generated in (8). At this point, you should have 3 x 5-point standard curves for 15 total tubes.
11. Add 200 µL from each tube to a well in your microplate, so that B1, B2, and B3 hold the triplicate samples made in step 8; C1, C2, and C3 hold the triplicates made in (9), etc.

12. With a volunteer’s help, measure the absorbance of each well at 628 nm. Copy the values into your notebook and generate a standard curve on Excel, plotting absorbance vs concentration. Add a linear trendline and use Excel to compute the trendline fit equation and $R^2$.

*What is the significance of the trendline equation? What is the significance of $R^2$? What value should $R^2$ be above in order to use the trendline?*

13. 2% (w/v) and 4% sodium alginate solutions have been prepared. In new epitubes, add 10 µL of stock dye to 990 µL of each sodium alginate stock.
14. Using a wide-bore pipette tip on the P1000, slowly and gently extrude 100 µL of coloured sodium alginate into 5 mL of $CaCl_2$ (10% w/v).

*What happens to the alginate?*

15. Repeat step 14 5x times.
16. Let the alginate sit for 15 min, then gently drain off the liquid. Add each bead to a new epitube containing 1 mL of water. Do this with both 2% and 4% solutions of alginate.
17. At 5, 10, 20, 30, and 60 minutes post-addition, take 200 µL from one tube and add it to a new well of the microplate. Once you have completed the time course, measure the absorbance of all wells at 628 nm.

**Copy these values into your notebook.**

**Calculate the amount of dye (in mol) released into each tube (remember you are only sampling 200 µL of a 1000 µL volume). Generate release curves (amount in mol released vs. time in minutes), in triplicate, of both 2% and 4% alginate solutions. What trends do you observe?**

S1 Appendix: Sample teaching materials for one term of IBBME Discovery

#### Chemistry request for proposal

The Institute of Biomaterials and Biomedical Engineering (University of Toronto) is working to improve the lives of patients around the world. Currently in Canada there are over 1.6 million people with cardiovascular disease, and cardiac health is one of the leading causes of death in the developed world. These patients suffer from a significant reduction in their quality of life as reduction in heart function makes everyday life difficult. Although healthcare continues to advance treatment, there is a need for continued focus on preventing, identifying and treating this prominent health problem. We are looking for the next generation of great minds to help us improve the lives of these people around the world!

Metoprolol is a beta-blocker that is frequently used to treat high blood pressure and tachycardia (chronically elevated heart rate). It can be taken by pill or intravenously but is a “dirty drug” in that it has numerous side effects, including depression, insomnia, fatigue, dangerously low heart rate, or respiratory depression. Part of this toxicity can be avoided by tightly regulating the concentration of the drug within the body. **We are working on bringing a new metaprolol treatment to market in either pill or implant form, but need chemical engineering expertise to optimize the release of the drug within safe limits. We want you to design a metoprolol-releasing hydrogel and quantify its rate of release to fall within safe limits.**

We will be collecting proposals from multiple teams, and will consider all options at a collective symposium. Please submit your project proposal using the RFP template provided. We have attached a list of the materials available in our facilities, so please take a look at these as you build your experimental plan. These will be reviewed by IBBME representatives who will meet with you to finalize the experimental details. You will then utilize our facilities to carry out your project and deliver a final presentation in poster format (template provided). We are looking forward to seeing the innovative solutions you will develop!

**Notes:**

**Be aware of experimental constraints**

You will have access to calcium and alginate solutions, food dye, materials to cast gels, microplates and spectrophotometers, and potentially other chemicals or materials available upon request.

**Do your research**

- How can we modify hydrogels to change delivery of a drug?
- What changes to current treatments would patients benefit from?

**Engineering is an iterative and creative process**

Some ideas you have might not be directly in line with the precise problem posed. There are plenty of issues in healthcare today, and given appropriate resources, you can adapt the project to your interest!

**If your proposal is approved, be prepared to make it happen!**

- Have a step-by-step protocol ready
- Be prepared to troubleshoot difficulties as they arrive
- Practice with Excel (and the sample worksheet) might be very useful!

### Physics

#### Physics project outline

**Motivation:** Currently in Canada there are over 1.6 million people with cardiovascular disease, and cardiac health is one of the leading causes of death in the developed world. These patients suffer from a significant reduction in their quality of life as reduction in heart function makes everyday life difficult. Although healthcare continues to advance treatment, there is a need for continued focus on preventing, identifying and treating this prominent health problem.

Improved resources for patients with cardiovascular disease to monitor their conditions would allow them to track parameters of interest, and schedule appointments or alert emergency services as needed if key metrics of cardiovascular performance change. A portable infrared plethysmograph capable of tracking pulsatile blood flow can be easily built using simple Arduino-compatible components.

**Research question:** Can we build a cost-effective infrared pulse sensor device to monitor cardiac function?

**Rationale:** Biomedical instrumentation is necessary to monitor and diagnose medical conditions. Students will learn the scientific basis of biological signals and the technical principles involved in detecting and measuring them. Close monitoring of a condition is also necessary for optimized intervention.

This project would also complement the “Waves and Sound” and “Electricity and Magnetism” units from the grade 11 and 12 physics curricula:

**Skill-based activity:** Several small tasks will teach the students some of the following concepts:

- Fundamentals of circuits: basic electronic components, units and concepts associated with electricity, and voltage/current laws.

- Signal Processing: signal filtering and amplification
- Coding: Writing code to calculate and display heart rate

The skill-based learning activity will involve a guided-session where students are introduced to the Arduino, a popular microcontroller, which is effectively a very simple computer. Students will follow steps to setup the Arduino, learn about its various components and features and how to upload code to the Arduino. At the end of the skill laboratory, students will have created a simple circuit interfaced with the Arduino that allows them to control the blink frequency of an LED by adjusting a potentiometer (a variable resistor device). This will teach skills including basic circuit design, collecting signals into the Arduino (a process known as analog-to-digital conversion) and how to send signals from the Arduino to an LED.

**Project:** Students will create a prototype of an IR pulse sensor/plethysmography device for measuring pulsatile blood flow in an extremity. This device and the process of creating this device will expose students to biomedical device design and development. By combining the scientific principles of cardiac physiology and engineering students will see first hand how scientific research can be translated into an engineering device that can be used to improve patients' quality of life.

Required equipment

- Laptops (for coding and 3D design)
- Arduino Microcontrollers
- Prototyping boards// breadboards
- Resistors + Capacitors
- Pulse Sensors (Contains LED, light sensor, and analog filter + amplifier)
- 3D Printers

**Expected outcomes:** At the end of this module, students will be able to:

- Understand basic electronic components and be able to plan and assemble rudimentary electronic circuits
- Obtain, modify, and run basic Arduino code
- Understand basic data acquisition and filtering processes
- Understand principles of 3-dimensional CAD and additive manufacturing

**Relevant literature topics:** Custom protocols and presentations will be provided. Readings on equilibrium, conservation of matter, etc. in their own textbooks will be helpful for review.

#### Physics skill lab

Purpose

This lab will introduce several important microcontroller concepts and features. First, it introduces the concepts of inputs and outputs. Inputs can allow the Arduino to connect to sensors, which detect variables from its environment (such as temperature) or allow a user to interact with it through buttons or a potentiometer. On the other hand, outputs allow the microcontroller to control other components or devices in order to display information back to the user or run a system. In this lab, we will see how programming the microcontroller allows us to control the relationship between those inputs and outputs.

Materials

- PC + Arduino IDE
- Arduino microcontroller + USB cable
- Breadboard
- 10kΩ potentiometer
- LED
- Jumper Wires

Procedure

*Part 1: Hook up an LED and make it flash at the rate of 1Hz:*

1. Connect the positive rail on the breadboard to +5V on the Arduino and the ground rail to any GND pin on the Arduino.
2. On the breadboard: Following the diagram in *Figure 1*, connect the digital output pin D13, the 100Ω resistor, the LED, and the ground rail *in series*. Note that the LED has polarity [a].

3. Connect your Arduino to a PC using the provided USB cable

4. In the Arduino <u>IDE</u> [b] open the Blink example code and set the blink rate by changing the number in the delay() function.

5. Compile your code by clicking this button in the Arduino IDE: 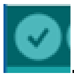. Compiling your code checks your code for errors and tells you if there are any problems.

6. Next, upload your code to the Arduino by clicking this button: 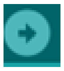

7. Voila, you're done! Enjoy endless hours of watching your LED blink!

[a] *Remember, as a result of polarity you can only insert the LED into the circuit in one way. Do some basic research on this.*

[b] *An IDE or "integrated development environment" is a program that you can use to write and then send code to the Arduino*

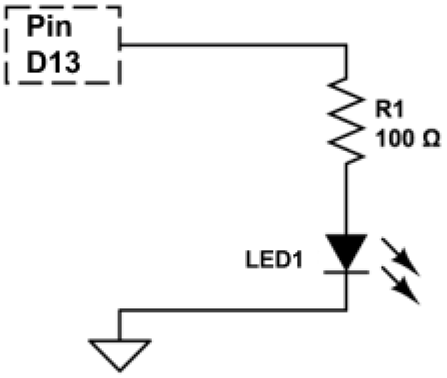

*Figure 1: The LED should be hooked up in series with a resistor. Remember to consider the connectivity of the circuit, not the geometry!*

*Part 2: Controlling the rate of an LED using a potentiometer:*

1. Do not change the circuit from Part 1!
2. On a different part of the breadboard, connect the circuit shown in *Figure 2*: connect the **positive rail** to one end of the potentiometer, the **ground** rail to the other, and connect analog input pin **A0** to the wiper or center pin on the potentiometer

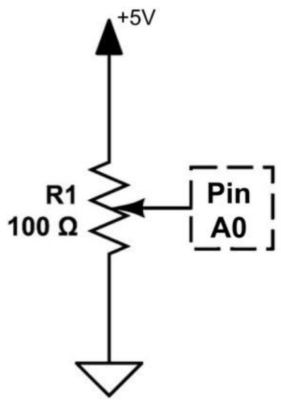

*Figure 2: The potentiometer circuit to control the LED blink rate*

3. Upload the controlBlink.ino file to the Arduino.
4. Now move the potentiometer and observe how it changes the blinking rate of the LED. Pretty cool eh!?

#### Physics request for proposal

The Institute of Biomaterials and Biomedical Engineering (University of Toronto) is working to improve the lives of patients around the world. Currently in Canada there are over 1.6 million people with cardiovascular disease, and cardiac health is one of the leading causes of death in the developed world. These patients suffer from a significant reduction in their quality of life as reduction in heart function makes everyday life difficult. Although healthcare continues to advance treatment, there is a need for continued focus on preventing, identifying and treating this prominent health problem. We are looking for the next generation of great minds to help us improve the lives of these people around the world!

Early detection of abnormal heart rhythms is important in order to provide timely and appropriate treatment for cardiac conditions. There are two main techniques that can be used to monitor a person's heart rhythm. These include measuring the electrical activity of the heart (an *electrocardiogram*) or detecting changes in the amount of oxygenated blood within a tissue region (e.g. a fingertip) based on the amount of infrared (IR) light that is reflected or transmitted (known as *IR plethysmography*). This works because the oxygenated blood contains higher levels of oxy-hemoglobin (hemoglobin is a compound in red blood cells which transports oxygen around the body), which reflects more infrared light than deoxygenated blood. **We are looking for a cost-effective device to measure a patient's heart rate. The design should be able to be easily prototyped and demonstrate the conceptual design of the device. The device should be comfortable, easy-to-use, and portable. It should be able to indicate the user's heart rate and alert them if there are measured abnormalities.**

We will be collecting proposals from multiple teams and will consider all options at a collective symposium. Please submit your project proposal using the RFP template provided. We have attached a list of the materials available in our facilities, so please take a look at these as you build your experimental plan. These will be reviewed by IBBME representatives who will meet with you to finalize the experimental details. You will then utilize our facilities to carry out your project and deliver a final presentation in poster format (template provided). We are looking forward to seeing the innovative solutions you will develop!

**Notes:**

**Be aware of experimental constraints**

*Time:* ~4 hours per visit, 2 visits

*Materials & Equipment:* See attached inventory list

**Do your research**

- Why is it important to monitor heart rate?
- What are some techniques that can be used to measure heart-rate?
- Compare the two most common techniques. What are their advantages and disadvantages?
- What is IR plethysmography (photoplethysmography)? How does it work?
- What are normal ranges for heart-rate?
- How can the device give feedback to the user?

**If your proposal is approved, be prepared to make it happen!**

Have detailed protocols ready and be aware of safety considerations around fabrication and electronics.

### Research Proposal Template

The following is a condensed version of the proposal template students complete in response to their subject specific RFP each term. Blue text is descriptors that guides student writing into specific categories, which is removed prior to submission.

##### TEMPLATE INTRODUCTION

*Use this template to respond to the "Request for Proposal" from IBBME Discovery, thereby developing a proposal for executing your experimental plan. The document should include: an introduction to the subject area and background research; scope and specific objectives; a detailed experimental plan including required materials and stepwise procedure as appropriate; breakdown of project management including a breakdown of team member responsibilities; and a summary of your expected outcomes. Throughout this template, italicized blue text is a descriptor of what information must be provided in a particular section, and therefore must be replaced with your own information. Remove all sections that include blue text from your submitted version, including this template introduction section.*

##### BACKGROUND RESEARCH & INFORMATION

*<A brief overview and background information>*

*A summary of the initial research for this project*

- *This is not a copy and paste of the background essay assignment*
    - *Include any information from the initial research that is relevant to the experiments you will be conducting*
- *Try to answer the following questions in this section:*
    - *What is the problem?*
    - *What effect does this problem have on society? Why is it important to solve?*
    - *What general approach will you be taking to solve this problem?*

- *Finally, what are you planning to design/test and how will this contribute to solving the problem?*

**SCOPE OF WORK**

*<A short introduction about the project and what testing will be performed>*

*What will you be designing in this project?*

*What need will this meet?*

*What is the general approach to test/evaluate your design?*

*What is the end goal?*

**SPECIFIC OBJECTIVES**

*<This Section describes the project structure, outlining which areas of focus you choose and why.>*

*What specific conditions will you be testing?*

*What specific tests will be carried out?*

*What tools will you use for performing these tests?*

**PROJECT REQUIREMENTS FOR ANALYSIS/EXPERIMENTS**

*<This is the core section in any research proposal. Each project requirement should be listed and described in specific detail. This section will outline your intended experiments. Think of this as being similar to materials and methodology for lab experiments.>*

**Materials**

*What do you need to create your design and carry out your experiments?*

**Procedure**

*<Describe the steps of your design and testing. Describe the main steps you plan to take to achieve the desired outcome. Provide a detailed list of what you will do in the experimental workspace>*

*What will you make?*

*What measurements will you take?*

*How will you analyze the data?*

*If you find that you need to make changes to your design, what changes will you make? What steps of your procedure will you perform again after?*

*Consider your variables – what will you change? What will you control?*

*What tests will you do?*

*What are your controls?*

*Examples of content to include:*

*Solutions you will prepare and respective dilutions*

*Preliminary design you have in mind and steps you will follow to fabricate*

*Experimental groups that will be considered.*

**RESPONSIBILITIES IN PROJECT MANAGEMENT**

*This section outlines when you will achieve each part of your proposed project and who is responsible for each step. Explain who is responsible for which deliverables. If a deliverable has more than one person assigned, detail each person's contributions. Indicate the planned timeline for accomplishing each task*

Deliverables & Parties Responsible

| Responsible Party | Date | Deliverable |
|---|---|---|

**EXPECTED OUTCOMES**

*<Summarize what you have proposed to do.>*

*What do you expect you will be able to demonstrate with this work?*

*How will you know you have succeeded?*

*Summarize briefly how this work can address the need you described in the background information.*

### Acknowledgements, usage, and adaptations

All Discovery materials are provided under the Creative Commons Attribution 4.0 (CC BY 4.0) license allowing the adaptation and usage of these materials for other educational purposes, provided that credit is given to the original authors and the Discovery program, and indication of any changes made is provided. Discovery does not endorse any external use or modification of these materials.

Discovery Biology materials were conceived and developed by Dr. Henrik Persson, Huntley Chang, Romario Regeenes, Cindy Yip, Janice Wong, Neal Callaghan, and Locke Davenport Huyer. Discovery Chemistry materials were conceived and developed by Neal Callaghan and Locke Davenport Huyer. Discovery Physics materials were conceived and developed by Daniel Smieja, Rami Saab, Neal Callaghan, and Locke Davenport Huyer.
