## Appendix 2 for "Enhancing senior high school student engagement and academic performance using an inclusive and scalable inquiry-based program"

### S2 Appendix: Entrance and exit student surveys for program assessment of IBBME Discovery

**Discovery Entrance Survey**

Name: ___________________________

1. In which course(s) are you enrolled: Physics Chemistry Biology

2. What grade are you in: 11 12

3. Have you participated in IBBME Discovery before? Yes No

   a. In what subject? (circle all that apply) Physics Chemistry Biology

   b. In which term? (circle all that apply)

   Fall 2016 Spring 2017 Fall 2017 Spring 2018 Fall 2018

4. How do you feel about taking science or science-focused courses in the future? (Circle one answer)

   a. I am likely to take science courses in the future.

   b. I am not likely to take science courses in the future

5. In your own words, what is biomedical engineering?

   ______________________________________________________________

6. Are you planning to graduate high school this year? Yes No

7. Did you apply to post-secondary institutions? Yes No

8. a. Where did you apply?

____________________________________________________________

____________________________________________________________

b. To what program(s) did you apply?

____________________________________________________________

____________________________________________________________

c. What factors are important in selecting a post-secondary destination (circle all that apply)

| | | | | |
|---|---|---|---|---|
| Program | Far from Home | Reputation | Social Environment | Personal Fit |
| Tuition Cost | School Size | Near Home | Campus Size | Athletics |

Available financial aid

Other: __________________________________________________

Additional Comments:

#### Discovery Exit Survey

Name: ___________________________

1. What project did you work on? Physics Chemistry Biology

2. What grade are you in? 11 12

3. What is one thing you learned from this project?

   ____________________________________________________________________________

   ____________________________________________________________________________

   ____________________________________________________________________________

4. Did you feel well-prepared for the project? Yes No

5. Did you feel challenged by the project? (Circle one number)

   *Yes I felt challenged academically* **5** **4** **3** **2** **1** *No I did not feel challenged academically*

6. Did this experience make you feel more comfortable with the idea of doing lab work in university or college?

   Yes No

7. What did you find to be the most difficult aspect to the project?

   ____________________________________________________________________________

   ____________________________________________________________________________

   ____________________________________________________________________________

8. Did you feel comfortable discussing your project in the Symposium setting? Yes No

9. How do you feel about taking science or science-focused courses in the future (Circle one answer)?

a. I am more likely to continue in science or engineering after IBBME Discovery.

b. I am less likely to continue in science or engineering after IBBME Discovery

c. IBBME Discovery had no influence on my future course selection.

10. Would you be interested in participating in a program like this again? Yes No

11. Have you participated in IBBME Discovery before? Yes No

a. In what subject (circle all that apply): Physics Chemistry Biology

b. In which term (circle all that apply):

Fall 2016 Spring 2017 Fall 2017 Spring 2018 Fall 2018

12. In your own words, what is biomedical engineering?

________________________________________________________________________

________________________________________________________________________

13. Are you planning to graduate high school this year? Yes No

14. Did you apply to post-secondary institutions? Yes No

15. a. Where did you apply?

______________________________________________________________

__________________________________________________________________________

b. To what program(s) did you apply?

__________________________________________________________________________

__________________________________________________________________________

c. What factors are important in selecting a post-secondary destination (circle all that apply)

| | | | | |
|---|---|---|---|---|
| Program | Far from Home | Reputation | Social Environment | Personal Fit |
| Tuition Cost | School Size | Near Home | Campus Size | Athletics |
| Available financial aid | | | | |

Other: __________________________________________________________________

Additional Comments:

*Thank you for your participation!*
