## Appendix 3 for "Enhancing senior high school student engagement and academic performance using an inclusive and scalable inquiry-based program"

### S3 Appendix: Mark breakdown for student assessment in *Discovery* programming

The following table provides a mark breakdown of the assessments and evaluations that comprise the *Discovery* cumulative assessment for participating students. All assessment was performed by participating educators for their respective students according to this template. It includes a mixture of written groupwork (Final Proposal, Progress Report, Poster), individual written work (Background Essay), and group presentation (Client Meeting, Poster Presentation). Percentage weight for each deliverable is divided to ensure that (1) there are both group and individual components of assessment and (2) students are assessed on both written and oral communication of their work.

*Table 1: Summary of mark breakdown for Discovery programming*

| **ITEM** | **DESCRIPTION OF ASSESSMENT** | **POINTS (/145)** | **PERCENTAGE OF GRADE** |
|---|---|---|---|
| **BACKGROUND ESSAY** | Relevant summary of research topic motivation | 20 | 13.8% |
| **CLIENT MEETING** | Student presentation skills in virtual client meeting that pitches proposed workflow in response to subject specific request for proposals | 5 | 3.4% |
| **FINAL PROPOSAL** | Group submission of planned workflow for the skill lab visits.<br>Assessment of workplan completeness, appropriate research question, hypothesis, and expected outcomes.<br>Description of groupwork breakdown clearly described | 35 | 24.1% |
| **PROGRESS REPORT** | Reflection on first skill lab visit outcomes, summary of progress to date<br>Updated plan of next steps to ensure research outcomes are complete at the end of next skill lab visit | 15 | 10.3% |
| **POSTER** | Assessment of poster content:<br>Research question and motivation clearly outlined<br>Clear communication of results<br>Reasonable conclusions drawn<br>Appropriate references to relevant literature | 40 | 27.6% |
| **POSTER PRESENTATION** | Assessment of each student's communication skills in poster presentation setting. Graded on ability to succinctly communicate research outcomes and answer relevant questions | 30 | 20.7% |
